# Relationship between X chromosome mosaicism, neuroanatomy and cognitive performance in females

**DOI:** 10.64898/2026.02.07.704538

**Authors:** Iliana I. Karipidis, Megan Klabunde, Tracy Jordan, Lindsay Chromik, S. M. Hadi Hosseini, Allan L. Reiss, David S. Hong

**Affiliations:** Center for Interdisciplinary Brain Sciences, Department of Psychiatry and Behavioral Sciences, School of Medicine, Stanford University, USA; Department of Child and Adolescent Psychiatry and Psychotherapy, Psychiatric University Hospital Zurich, University of Zurich, Switzerland; Neuroscience Center Zurich, University and ETH Zurich, Zurich, Switzerland; Centre for Brain Science, Psychology Department, University of Essex, United Kingdom

## Abstract

Females have two X chromosomes, one of which is inactivated early in development with specific regions and genes ‘escaping’ inactivation. Thus, X chromosome loss putatively results in decreased ‘dosage’ of X chromosome escapee and pseudoautosomal genes, impacting downstream pathways. Evidence from Turner syndrome indicates that X chromosome monosomy results in consistent neuroanatomical and cognitive phenotypes. However, it remains unclear whether mosaic karyotypes, with mixed proportions of 45X and 46XX cells, attenuate these phenotypes. We examined whether X chromosome mosaicism is predicted by neuroanatomical and cognitive features. Higher proportion of 46XX cells was significantly predicted by structural properties in somatosensory, motor, visual, and language brain areas, and by performance in visuospatial, fine-motor, and language tasks. Thus, mosaicism partially ‘rescues’ phenotypes linked to full 45X monosomy and may explain the role of the X chromosome not only across heterogeneous phenotypic expression in females, but also in sex differences observed in neuropsychiatric conditions.

## Introduction

The X chromosome is increasingly emerging as an important genetic driver in neurodevelopment and brain organization(1), yet has also been historically understudied due to analytic challenges in managing X-Y chromosomal imbalance. Recent establishment of large genetic and neuroimaging databases (2, 3) has facilitated unprecedented insights into the associations between genetic variability and complex neurobiological phenotypes at the population level in humans. At the same time, genetics-first approaches investigating specific groups of individuals with a shared etiology described by a well-defined set of genetic variations has enabled understanding of how genetic causes may result in related clinical phenotypes, as is the case in chromosomal aneuploidies (4). However, much remains to be delineated in understanding the pathways by which a genetic variation results in a neuroendophenotype, and an associated cognitive-behavioral outcome, given the inherent complexity within each of these biological domains.

One compelling mechanism that may shed important insights into these relationships is the concept of gene ‘dosage’, i.e., whether downstream biological products from a genetic variation additively increase the loading of a related phenotype (5). Dosage sensitivity has been well-established as a biological driver for pathogenicity in human neurodevelopmental disorders (6), as operationalized by copy number variation in genomic regions typically spanning multiple genes. This principle is also relevant at a larger order of magnitude, encompassing variation in whole chromosome number. The most extreme representation of this are chromosomal aneuploidies resulting in penetrant phenotypes, e.g. Trisomy 21, but is also relevant for mosaic karyotypes within aneuploid states.

Specific to X chromosome biology, the gene dosage model is a compelling framework to characterize sex chromosome aneuploidies (SCAs), such as Turner syndrome (TS; 45X), where chromosomal mosaicism commonly occurs (7). Aneuploidy in sex chromosomes is distinctive given their non-pathological dosage compensation. In females (46XX), most of one X chromosome is inactivated to compensate for far fewer genes on the Y chromosome in males (46XY) (8). This results in a monoallelic state for most X chromosome genes, excepting ‘escapee’, pseudoautosomal, and X-Y homolog genes (9), which are then functionally haploinsufficient in TS, relative to euploid cells. Within this context, TS is an aneuploidy defined by complete or partial monosomy of the X chromosome. Approximately half of individuals with TS carry the most common karyotype, 45X complete monosomy (10). The associated haploinsufficiency of undefined X chromosome genes are putatively linked to a well-established neurocognitive phenotype that is usually characterized by intact full-scale IQ, with deficits in visual-spatial abilities, fine motor skills, executive functions, and social-cognition (11–15). In parallel, monosomic TS (monoTS) is also associated with clear structural and functional neuroendophenotypes. Namely, TS is associated with reduced cortical volume (CV) in occipital and parietal brain areas, while CV of the superior temporal cortex has been systematically reported to be higher in TS compared to control individuals (16–22). Relatedly, comparison of CV in TS with Klinefelter syndrome (KS, 47XXY) has provided first evidence that these deviations in CV likely originate from gene dosage effects in the sex chromosomes (23).

Important for this investigative framework, approximately 17% of individuals with TS have a 45X/46XX mosaic karyotype, where a proportion of cells have one X chromosome (45X) and a complementary proportion have a typical 46XX karyotype (10). Critically, TS is the only human complete monosomy compatible with life, likely due to existing dosage compensation mechanisms for X-Y imbalance, as well as balance with autosomal dosage (24–26). This uniquely makes mosaic TS (mosTS) one of the only feasible frameworks in which to investigate specific dosage effects of X chromosome gene products on human phenotypes. Chimeric distribution of euploid and aneuploid states presumably results in attenuation of the expected phenotype associated with full monosomy (27). Validation of this framework would allow critical insights into the genotype-phenotype relationship in TS and how it is moderated by mosaicism – therefore, mosTS represents the only in vivo human model permitting identification of neurobiological and cognitive-behavioral domains that are sensitive to genetic ‘rescue’ from a known *monosomic* phenotype via X chromosome dosage (XCD) effects.

Indeed, early small-scale studies of TS with varied karyotypes suggest that clinical characteristics of TS may be attenuated by the mosaic karyotype (28, 29), such as mosTS likely being associated with slightly higher levels of intellectual ability than monosomic karyotypes (30) and more balanced IQ profiles (31). This is additionally supported by evidence that on average, individuals with 45X monosomy perform worse than individuals with mosaic 45X/46XX karyotypes on cognitive and motor tasks (31–33). While genetic studies to date demonstrate genes at different loci of the X chromosome are linked to the physiological and hormonal characteristics in TS (32), including seminal evidence that stature is at least partly dosage-sensitive due to copy number variation effects from the SHOX gene (34), dosage sensitivity of sex chromosome genes contributing to the neurodevelopment and neurocognitive profiles remain elusive, despite emerging evidence that the X chromosome has an outsized impact on brain development in particular, even in euploid populations (1).

To address this gap in knowledge, we directly examined 45X/46XX mosaicism effects on cognition and neuroanatomy through investigation of the relationship between gene dosage, neuropsychological measures, and brain structure in pre- and early pubertal girls with and without TS, providing the first predictive model of structural brain and cognitive measures based on percentage of X chromosome mosaicism. To do so, we implemented support vector regression, a well-established, supervised machine learning approach that uses a multidimensional regression model to perform predictions. We examined phenotypes by modeling a continuum of X chromosome ploidy, as operationalized by individuals with varying levels of X chromosome mosaicism without structural abnormalities (e.g. translocations, ring, deletions), individuals with 100% 45X karyotypes (i.e. monoTS), and individuals with 0% 45X mosaicism (i.e. euploid 46XX karyotypes). Using this approach, we 1) evaluated sensitivity to percentage of X chromosome mosaicism in neuropsychological performance, i.e. tested which scores across the domains of sensorimotor, visuospatial, language, memory, executive functions, and social perception can predict XCD, 2) identified X chromosome sensitive regions across CV, cortical thickness (CT), and cortical surface area (CSA), and 3) implemented a multimodal approach combining neuropsychological and brain structure measures.

## Results

### Neurocognitive performance predicts degree of X chromosome mosaicism

To investigate XCD effects on cognition, we selected 56 neurocognitive scores from an extensive neurocognitive test battery that was performed with pre- and early pubertal girls (Tanner stages 1 and 2) with varying degrees of mosaicism of the X chromosome (n = 14, 9.1±2.1y), monoTS (n = 15; 10.1±2.1y) and controls with euploid number of X chromosomes (n = 19, 9.4±1.5y). After principal component analysis, 11 principal components (PC) were entered into the SVR model to determine whether neurocognitive features could predict the percentage of X mosaicism ranging from 0 i.e., none of the tested cells was missing an X chromosome (46XX), to 100 i.e., all tested cells were missing the second X chromosome (45X).

After cross validation and non-parametric permutation testing, our multivariate model significantly predicted the degree of mosaicism in the test cases (rho = 0.33, P = 0.02, Fig 1A). Post-hoc feature weight analysis indicated that neurocognitive measures associated with sensorimotor processing, visual-spatial skills, verbal and language abilities, as well as attention and executive functions contributed the most to the prediction of X chromosome mosaicism (Fig 1B). Specifically, higher scores in visual-fine-motor tasks, including visuomotor precision, fine motor skills (WRAVMA Pegboard), fine motor coordination in imitating hand positions, drawing, spatial visualization and motor coordination (WISC Block Design), were predictive of a *lower* percentage of mosaicism. In addition, higher values in visual spatial measures, such as visual tracking and judging line orientation (NEPSY Arrows), categorical reasoning requiring figure-ground perception (WISC Picture Concepts), and processing speed in naming shapes requiring response inhibition (NEPSY Inhibition-Naming) also contributed more towards the prediction of increased proportion of 46XX cells.

**Figure 1.**
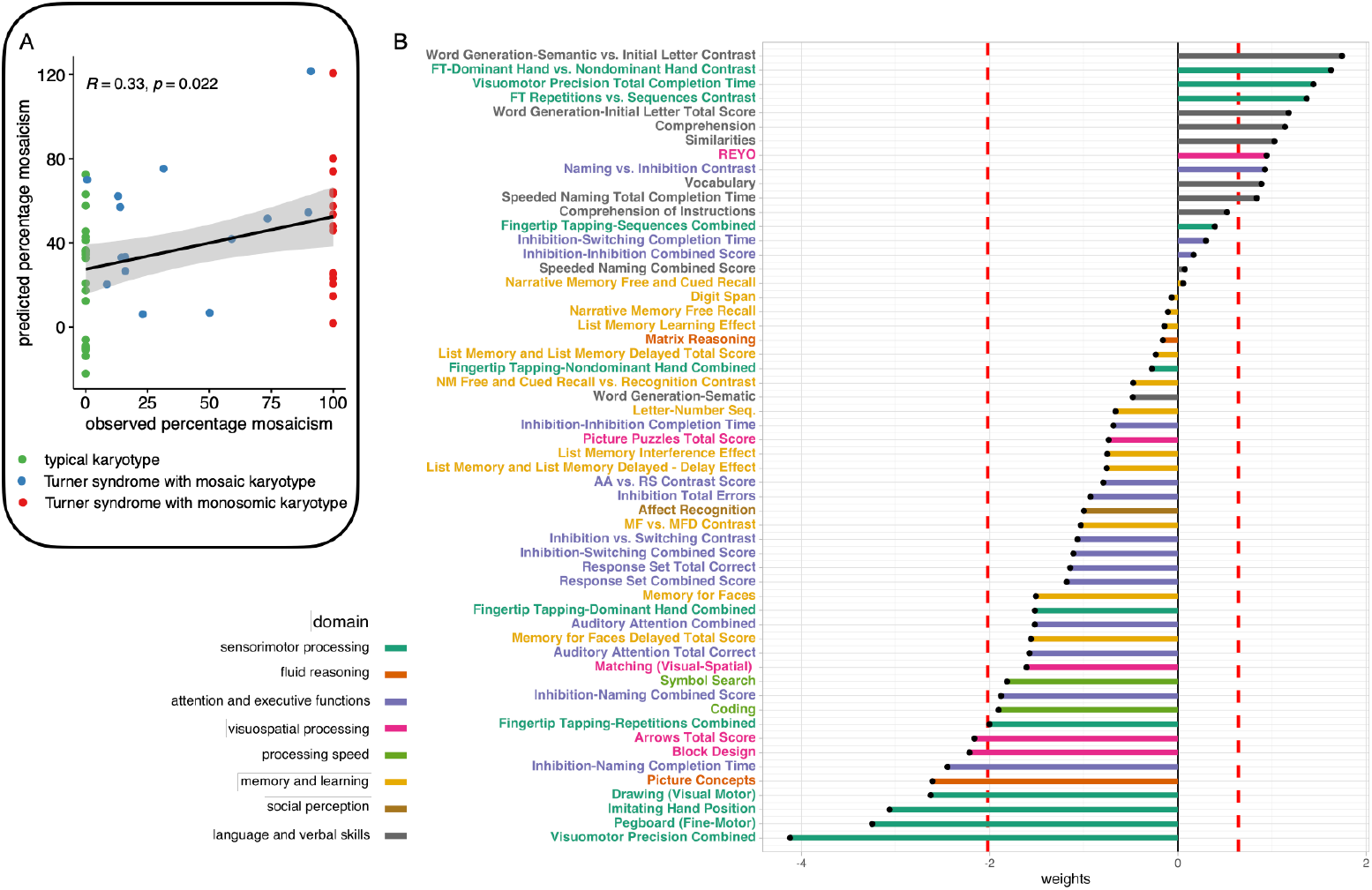
Support vector regression (SVR) results using neurocognitive measures. (A) Evaluation of cross-validated prediction accuracy using Spearman correlation. (B) Positive and negative weights of features contributing to the SVR model. Neurocognitive measures with positive weights indicate that higher performance is predictive of higher percentage mosaicism, while negative weights are more predictive of lower percentage mosaicism. Red dashed line marks 1 SD from the mean. Scores are derived from the Wechsler Intelligence Scale for Children (WISC-4), Rey-Osterrieth complex figure (REY-O), NEPSY: A Developmental Neuropsychological Assessment (NEPSY2), and the Wide Range Assessment of Visual Motor Abilities (WRAVMA).

We also found language-based domains were sensitive to XCD, including findings that scores in language comprehension (Comprehension, Similarities, and Vocabulary in WISC), verbal fluency (NEPSY Word Generation-Initial Letter), producing language effortlessly when organized categorically (Word Generation Semantic vs. Initial Letter), serial naming requiring inhibition (NEPSY Naming vs Inhibition Contrast), and word retrieval and producing verbal labels (NEPSY Speeded Naming) were predictive of a *higher* percentage of 45X mosaicism. In contrast, higher scores on primarily motor domains, including fine motor control of the dominant vs the non-dominant hand (NEPSY Fingertip Tapping), time needed to perform a manual motor task (NEPSY Visuomotor Precision), performance in motor programming compared to motor control (NEPSY Fingertip Tapping repetition vs sequences), and accurately and efficiently copying a complex figure (RCFT-REYO score), indicated a higher likelihood of X chromosome monosomy.

### X chromosome mosaicism and neuroanatomy

We then considered an SVR prediction model using neuroanatomical data derived from T1-weighted structural whole-brain MRI, operationalized through Freesurfer outputs of CV, CT, and CSA, resulting in 68, 68, and 70 cortical and subcortical parcels respectively. We applied the same procedure to construct three separate neuroanatomical SVR models to predict X chromosome mosaicism.

### Cortical Volume

The 68 volumetric measures were reduced to 14 PCs that were used to fit the SVR model. We observed that the cross-validated predicted percentage positively correlated with the true levels of 45X mosaicism (rho = .56; *p* < .001; Fig 2A). Post-hoc feature weight analysis indicated that the most important volumetric measures contributing to the prediction of 45X-chromosome mosaicism (1SD from the mean) were located in primary somatosensory regions, visual regions, the insulae, and frontal brain areas. Specifically, we noted increased CV bilaterally in the somatosensory and motor cortex, including the superior parietal lobes (SPL), postcentral gyri, and precunei was predictive of a lower percentage of X chromosome mosaicism. In addition, greater CV in the right pars opercularis and in visual regions, including bilaterally the lingual, fusiform, and pericalcarine gyri were also predictive of lower mosaicism. Conversely, greater CV bilaterally in the insula and in frontal regions, including rostral middle frontal gyrus, superior frontal gyrus, pars orbitalis, and the right lateral orbitofrontal cortex were predictive of a higher percentage of X chromosome mosaicism. Increased CV in the left posterior cingulate, left lateral occipital gyrus, and right precentral gyrus were also more likely to predict a higher percentage of mosaicism towards X chromosome monosomy.

**Figure 2.**
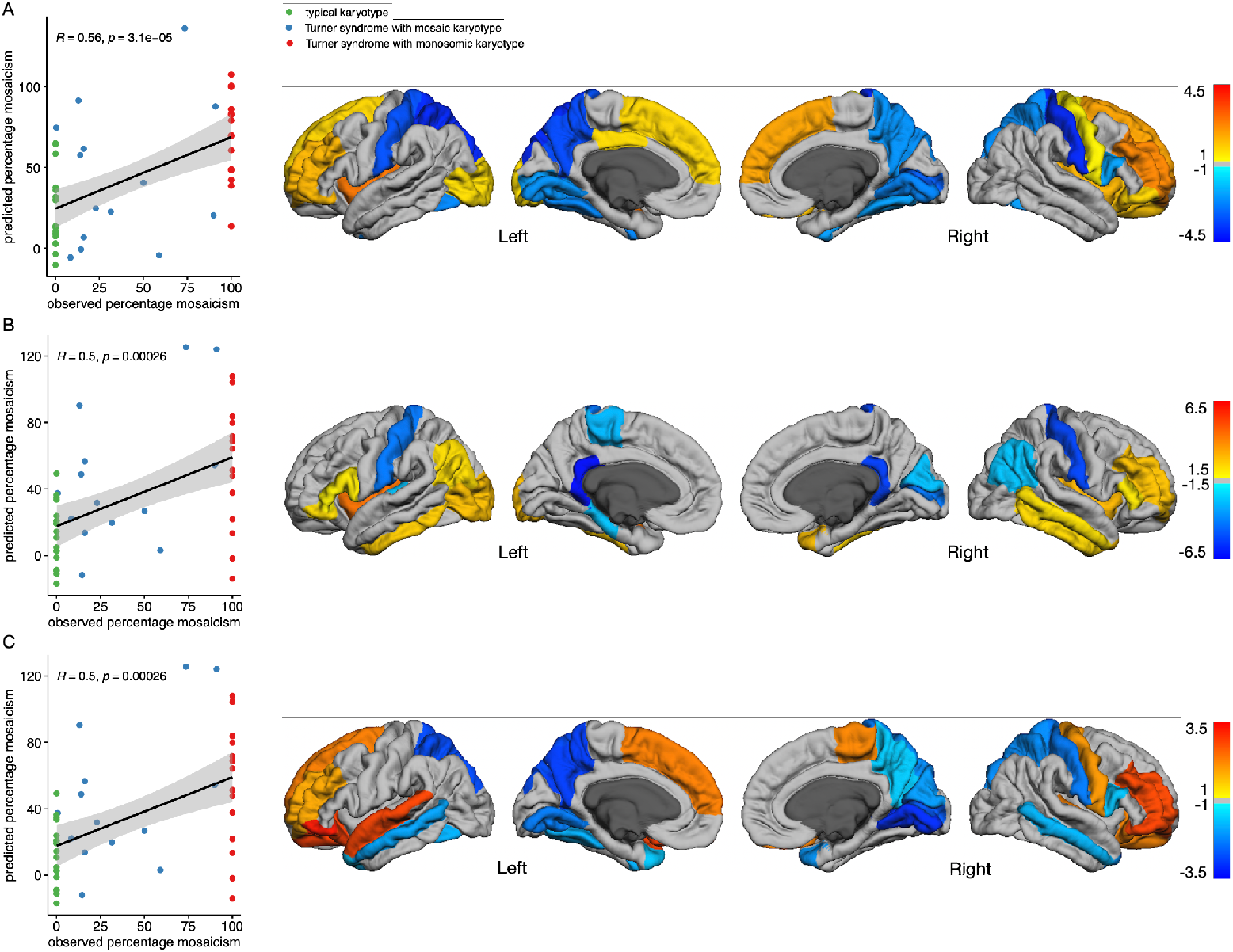
SVR results using structural brain measures. (A) Cortical volume (B) Cortical thickness (C) Surface area. Left: Evaluation of cross-validated prediction accuracy using Spearman correlations. Right: Visualization of features with the highest weights (>1SD from the mean).

### Cortical Thickness

We then reduced the 68 CT measurements to 14 PCs, which also significantly predicted X chromosome mosaicism percentage (rho = .52; *p* < .001; Fig 2B). Similar to the SVR CV model, increased CT in somatosensory regions was predictive of lower percentage mosaicism, while larger CT in the insula and frontal areas was predictive of a higher percentage of 45X mosaicism. We found that the most negative weights for this SVR model (indicating that increased CT is predictive of a lower percentage of 45X mosaicism) were bilaterally in the isthmi of the cingulate, extending to the left parahippocampal gyrus. Next to the postcentral gyri, greater CT in the left paracentral and right inferior parietal lobe were also predictive of a lower X chromosome mosaicism. Finally, increased CT in the left transverse temporal gyrus responsible for auditory processing, as well as parts of the right visual cortex, including the pericalcarine gyrus and cuneus were also more likely to predict lower mosaicism. We observed that the highest weights for the CT SVR features were found for the insulae, suggesting that increased CT was predictive of a higher percentage of 45X mosaicism. Additionally, increased CT in regions related to object recognition and language skills, including the left lateral occipital gyrus, left inferior parietal lobe, bilateral inferior temporal gyrus, right temporal pole, and right middle temporal gyrus were also found to be predictive of higher mosaicism. Lastly, increased CT in several frontal regions, such as the right rostral middle frontal gyrus, left pars orbitalis, left pars opercularis, and bilaterally pars triangularis, also predicted a higher percentage of 45X mosaicism.

### Surface Area

For the CSA SVR model, 70 Freesurfer measures were reduced to 14 PCs and again significantly predicted percentage mosaicism (rho = .50; *p* < .001; Fig 2C). Corresponding to the CV results, we observed that increased CSA in visual and somatosensory regions was predictive of low mosaicism, while increased CSA in frontal areas was predictive of high mosaicism. The lowest weights for CSA SVR features were found in visual regions, such as the lingual gyri, right pericalcarine cortex and right cuneus, as well as in parietal regions, including bilaterally the precuneus and superior parietal lobe, as well as the right postcentral somatosensory areas. In addition, we found that increased CSA in the temporal poles, middle temporal gyri, left fusiform gyrus, and right pars opercularis were predictive of lower percentage of mosaicism.

We found the highest weights for CSA SVR features in the pars orbitalis, a key region in language processing. Greater CSA in frontal regions, such as bilaterally in lateral orbitofrontal cortex, rostral middle frontal gyrus and left superior frontal gyrus, as well as in some right-hemispheric parietal regions, the paracentral and precentral gyri, was also predictive of a higher percentage of X chromosome mosaicism. Lastly, in line with previous findings in TS, we demonstrate that increased CSA in primary auditory regions, the left superior temporal gyrus and right transverse temporal gyrus, contributed to the prediction of increased proportion of 45X mosaicism.

### Joint model with neurocognitive measures and cortical volume

As we found that of the neuroanatomical measures used for SVR, CV provided the best fit and resulted in the highest prediction accuracy correlation, we then combined CV measures alongside all neurocognitive scores to build a joint SVR model comprised of 124 features, summarized in 13 PCs. We found a significant prediction of percentage 45X mosaicism with a high prediction correlation value that exceeded the SVR predictions using neurocognitive and volumetric measures in isolation (rho = .67; *p* < .001; Fig 3A). This finding suggests that neurocognitive and CV measures contribute to the prediction of 45X mosaicism in a complementary way.

**Figure 3.**
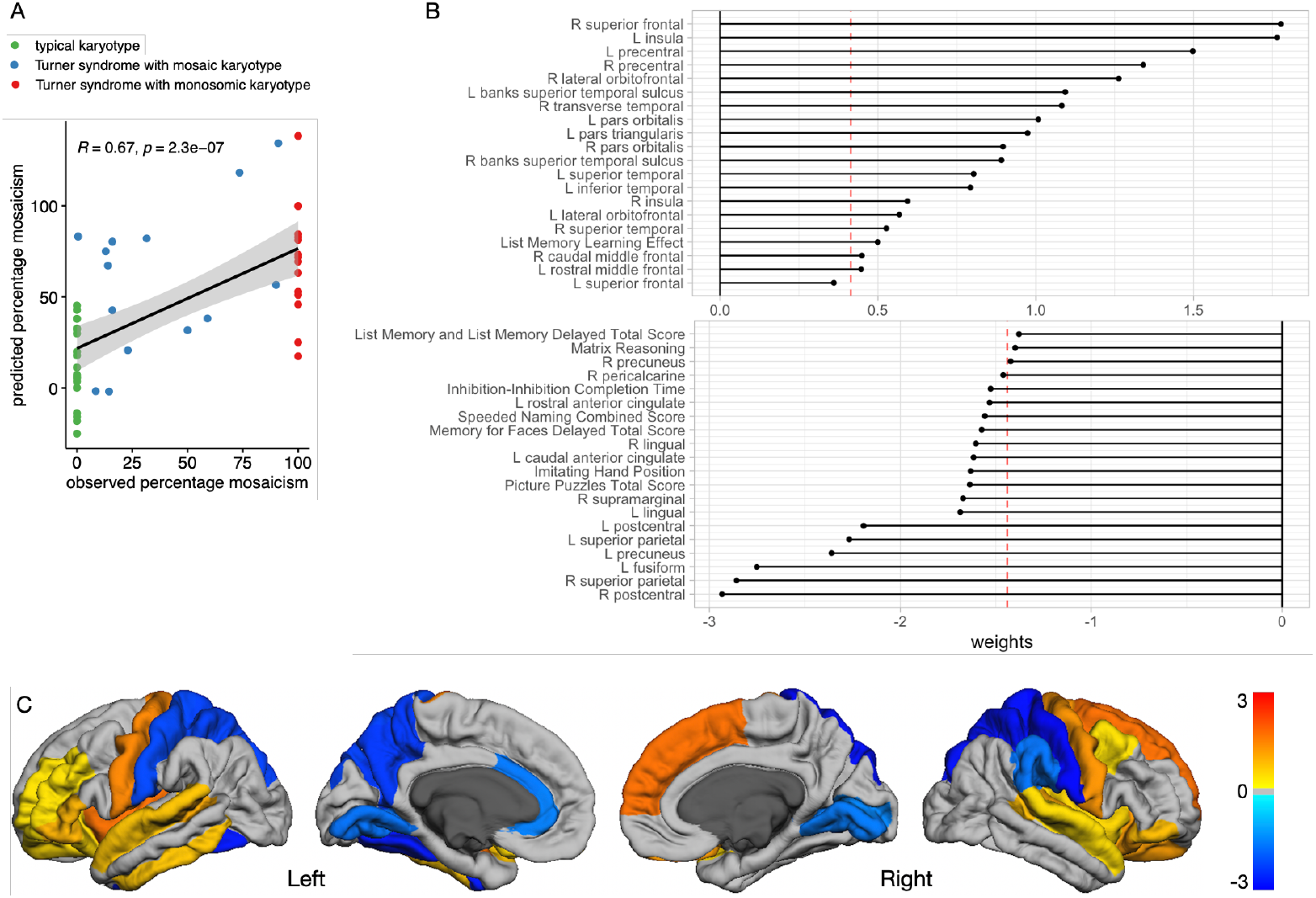
SVR results using and cortical volume and neurocognitive measures. (A) Evaluation of cross-validated prediction accuracy using Spearman correlation. (B) Square weights of the 40 features with the highest positive and negative contribution to the SVR model. Red dashed line marks 1 SD from the mean. (C) Visualization of cortical volume features with the highest square weights (>1SD from the mean).

The joint model confirmed that the lowest weights for volumetric and neurocognitive SVR features were found in somatosensory and visual domains. Increased CV in postcentral gyri and superior parietal lobes, as well as in other parietal regions, such as the left precuneus and right supramarginal gyrus, were predictive of a lower percentage of 45X mosaicism. Accordingly, neurocognitive scores associated with motor coordination i.e., imitating hand position also showed a very low feature weight. Additionally, we identified that greater CV in visual regions, including bilaterally in the lingual gyrus, left fusiform gyrus, and right pericalcarine gyrus, as well as the neurocognitive measure of visual perception (NEPSY Picture Puzzles) and learning (NEPSY Memory for Faces-Delayed) were also predictive of lower proportion of mosaicism. Lastly, the combined model also revealed that high scores in tests requiring impulse inhibition (Speeded Naming, Inhibition-Completion Time) and increased CV in the left anterior cingulate, known to be involved in impulse control, predicted a lower percentage of mosaicism.

In line with the SVR using only the volumetric data, the joint model showed that increased CV bilaterally in the insula, somatomotor areas (precentral gyri), and in frontal regions, including the right superior frontal gyrus, lateral OFC, pars orbitalis, left pars triangularis, right caudal middle frontal gyrus, and left rostral middle frontal gyrus were predictive of a higher percentage of 45X mosaicism. Additionally, the combined model revealed that increased CV in auditory regions, such as bilateral STS, bilateral STG, and the right transverse temporal gyrus (Heschl’s gyrus) predicted higher X chromosome mosaicism. The neurocognitive score with the greatest positive weight was associated with auditory processing ability as measured by performance in memorizing verbal information (NEPSY List Memory-Learning Effect). Lastly increased CV in the left inferior temporal gyrus was found to contribute to the prediction of higher percent mosaicism.

## Discussion

Here, we demonstrate that the degree of X chromosome mosaicism can be predicted by neurocognitive, CV, CT, and CSA measures. In one of the first studies to model a continuum of X chromosome ploidy through inclusion of mosaic karyotypes, we show a distribution of ‘intermediate’ phenotypes between patterns characteristic of 46XX and 45X brain anatomy (35), implicating a dosage-related genotype-phenotype relationship between X chromosome loss and neural and cognitive domains. Notably, using SVR we found that lower scores in visual-spatial and visual-fine-motor skills, and higher scores in verbal and language domains, predicted higher levels of 45X mosaicism, i.e. a greater proportion of detected 45X cells. Regarding brain-based measures, higher levels of X chromosome mosaicism were significantly predicted by lower CV, CT, and CSA in somatosensory, motor, and visual regions, and by higher CV, CT, and CSA measures in frontal regions, language areas, and the insula. The SVR model combining all neurocognitive and CV features corroborated these findings and additionally revealed that higher values in the domain of impulse inhibition and control explained lower percentage of X chromosome mosaicism, as did higher levels of motor coordination and CV in somatosensory regions.

The results revealed that cognitive features reflecting visual-fine-motor skills and visual-spatial abilities provided the strongest contribution towards predicting percentage of X ploidy. We also found significant contributions towards higher levels of X chromosome mosaicism for several features related to language and verbal skills. This is in line with previously described cognitive phenotypes in monoTS (36) and suggests that deficits in the visual-spatial domain are particularly attuned with higher X chromosome mosaicism, as are higher language and verbal skills. This is consistent with findings of deficits in visuospatial skills in monoTS (37), which clinically represents one of the most common neuropsychological impairments associated with TS (36). Additionally, difficulties in visuospatial processing in combination with spatiotemporal processing deficits lead to impairments in numerical cognition, likely underlying increased prevalence of dyscalculia among individuals with TS (38, 39). Relatedly, numerical cognition and visuospatial ability have been postulated to be influenced by dosage-sensitive X-linked genes (40), emphasizing the link between XCD and the broader phenotype associated with X chromosome loss. Notably, however, individuals with TS may be able to compensate for visuospatial deficits to some extent by relying on verbal skills during arithmetic processing (41). TS is therefore often characterized by an imbalance between verbal IQ and performance IQ, with individuals with TS usually scoring higher in verbal IQ than in performance IQ (30, 42). Motor deficits are also found in TS and are suggested to occur regardless of whether they are also contributed to by visual-spatial abilities.

The SVR model based on CV provides novel evidence that increasing levels of X chromosome mosaicism can be explained by lower CV in primary somatosensory and visual regions, and by higher CV in frontal areas and bilaterally in the insula. Results based on CT and CSA showed very similar effects, primarily confirming the predictive value of all three neuroanatomical components, which is not surprising given that CT and CSA are subcomponents of CV. However, we also identified unique contributions of regional CT and CSA measures in predicting X chromosome mosaicism, which were not evident when considering CV measures alone. Higher CT and CSA in temporal language areas was predictive of higher levels of mosaicism, which is in line with accounts reporting intact or even pronounced verbal and language skills in TS (43). Additionally, we found that lower CT in the right IPL and the isthmus of the cingulate cortex explained higher levels of X chromosome mosaicism. Given that the right IPL is implicated in mathematical cognition, lower CT in the right IPL with increasing degree of X chromosome mosaicism could be associated with difficulties in arithmetic cognition and the approximately 50% prevalence of dyscalculia among individuals with monoTS and mosTS (44).

Our findings demonstrate that individuals with mosaic karyotypes fall within the bounds defined on either end by ‘complete’ 45X monosomy or 46XX euploid karyotypes. Further, predicted scores are distributed in linear fashion, suggesting that effects of mosaicism are accumulative – or in the reverse view, the extent of mosaicism attenuates at least part of the neurocognitive and neuroanatomical phenotype linked with monoTS. This raises important implications for a dosage-based model of X chromosome effects for several reasons. Firstly, specific to the cognitive and neuroanatomical domains identified in the SVR, increasing presence of a second X chromosome, and related downstream gene products, putatively mediate haplosufficiency effects that drive previously observed X monosomic phenotypes. Given the broader understanding of X chromosome biology where dosage effects are feasible since X inactivation negates dosage imbalance between individuals with one X chromosome (45X) vs. two X chromosomes (46XX), these data provide further support that X chromosome genes, specifically, pseudoautosomal regions (PAR), escapee genes and X-Y homologs, are compelling candidates linked to visual spatial, motor, and to some degree verbal and language phenotypes. This framework is further bolstered by prior literature in boys with KS (47XXY), where increased XCD in a polyploidy model is correspondingly associated with improved visual spatial capacities and a protection from processing speed and executive functioning difficulties (23, 45). Indeed, when individuals with monoTS, KS and 46XX and XY karyotypes were compared together, similar neurocognitive and neuroanatomical regions were identified as in this study, suggesting the linear association extends beyond varying degrees of X chromosome loss and into excess X chromosome material as well.

Secondly, these findings provide important insight into sex differences even in individuals with euploid karyotypes. While these findings are presented in the context of aneuploidy, the broader relevance of an accumulative gene dosage approach to X chromosome genetic material is relevant to understanding sex-specific drivers in the general population. Despite the small sample size, our findings across a spectrum of X chromosome loss remarkably overlap with results derived from the UK Biobank, which identified many of the same neuroanatomical regions that differed between 46XY males and 46XX females (46). Those results demonstrated larger CV for males in the insula, orbitofrontal and occipitotemporal regions, and relatively larger CV for females in superior parietal, postcentral, and middle/inferior frontal areas among others. The regional neuroanatomical findings in our sample based on XCD correspond to these sex differences. In particular, brain regions with higher CV or CT in females than in males (46) were also associated with reduced X chromosome mosaicism in our analysis (increased XCD in left superior parietal gyrus, postcentral gyrus bilaterally, isthmus of cingulate gyrus, right pericalcarine gyrus), while some brain regions with higher CV or CSA in males compared to females were associated with increased X chromosome mosaicism (reduced XCD in right insula, left superior temporal gyrus, pars orbitalis bilaterally). These parallel results provide converging evidence for an X chromosome haploinsufficiency genetic framework, wherein ‘dosage-related’ loss of the second X chromosome as modeled in our mosTS data, approximates haplosufficiency of X chromosome escapee genes in 46XY males compared to 46XX females, at least for these cognitive and neuroanatomical domains. Notably, we also found that some regions did not overlap with UKBB data in adults (e.g. differences in CV for XY>XX in the fusiform and orbitofrontal gyrus). This may be in part related to developmental effects given our sample of early/pre-pubertal girls, which would be consistent with earlier findings of changing brain morphology driven by puberty (47, 48). Alternatively, non-overlapping regions may also indicate differing genetic etiologies – as opposed to sex differences in euploid populations, the regions identified here may reflect the impact of PAR genes and other X-Y homologs outside of the PAR (ZFY/ZFX and RPS4Y/RPS4X, amelogenein; and a block of sequences in Xp22.3-pter that includes the *STS, KAL*, and *GS1* genes showing homology to proximal Yq). Several of these regions have been implicated in sex differences and are particularly pronounced in SCAs.

A critical insight from this work is the potential role of mosaicism in observed inter-individual variation in neurocognitive and neuroanatomical phenotypes associated with TS. Our findings and prior TS literature show that despite shared genetic etiology, there is considerable variance in the penetrance of these genetic effects in TS, where even individuals with a 45X monosomic phenotype demonstrate substantially varying performance on implicated cognitive domains, including visual-spatial and motor performance. The findings here provide support for a mechanism in which mosaicism may explain at least part of this variation. One reason for this is that observed genotypes are based on a limited number of cells counted from peripheral blood. Within TS literature there is building evidence for cryptic mosaicism – namely, the concept that even individuals with a presumed 45X monosomic karyotypes have undetected cell lines with 46XX (58). Data from the UK Biobank suggests that the prevalence of mosaic 45X/46XX karyotype among adult women may be as high as 86% of all TS cases with many individuals remaining undiagnosed due to an unremarkable phenotype (49). Some findings even suggest that pure X chromosome monosomy may not be viable with life (50, 51). Thus, while we have demonstrated XCD effects in neuroendophenotypes of individuals with explicit mosaic karyotypes, this premise could hold true even with monoTS, providing additional support for degree of mosaicism representing a relevant biomarker in the clinical care of individuals with TS.

Our study should also be considered in the context of several limitations, including the relatively small, though carefully sampled, cohort of data included in this study. As diagnostic sophistication in identifying individuals with even limited mosaicism emerges, e.g. noninvasive prenatal testing, larger studies may reveal more refined associations than the ones demonstrated here. Further, these analyses were conducted through the lens of genetic effects driven by the X chromosome, however, it is difficult to fully segregate parallel hormonal effects related to estrogen differences between these cohorts. While this is partially mitigated by restriction of the sample to pre- and early pubertal girls, future studies should examine the role of sex steroid differences across developmental stages to further delineate the relationship between genetic and hormonal effects.

Lastly, it is also important to consider that tissue-specific and somatic mosaicism across organ systems may further contribute to interindividual variation in observed phenotypes. Evidence here indicates that at least at the level of neuroanatomy, mosaicism in peripheral blood mononuclear cells effectively correlates to variance in specific neuroanatomical domains. However, to the extent possible in pediatric populations, future studies should extend these investigations into central nervous system-derived cell types for broader understanding.

Despite these limitations, our study provides evidence that neurocognitive and structural MRI data can significantly predict degree of X chromosome ploidy, using the only dataset of well-characterized 45X/46XX mosTS to date. We demonstrate a linear distribution of X chromosome ‘gene dosage’ effects on neuroanatomical and behavioral phenotypes. We also surprisingly found that many of these regional brain differences were similarly identified as sex differences in substantially larger neuroimaging data sets between euploid males and females. The notable recapitulation of male-female sex differences in female-only cohort with varying degrees of X chromosome material, provides additional support for a putative dosage effect of X chromosome genes, highlighting the importance of further investigation into the role of X chromosome mosaicism, which may be much more prevalent that previously known, and underscores critical gaps in the prior evidence base in which sex chromosomes are typically omitted in genetic analyses (52). Future work may focus on X chromosome genes as potential therapeutic targets, as well as incorporation into clinical algorithms by which to guide neuropsychological functioning in youth with TS and to prioritize treatment approaches and prevention strategies in this population.

## Materials and Methods

### Participants

N=48 pre- and early pubertal girls (Tanner stages 1 and 2) with average or above average IQ and varying degrees of mosaicism of the X chromosome were recruited. The sample included 14 girls with mosaic TS (mean age 9.1±2.1y), 15 girls with monosomic TS (mean age 10.1±2.1y) and 19 age- and IQ-matched controls with euploid number of X chromosomes (mean age 9.4±1.5y). Participants with TS were recruited by local and national chapters of the Turner Syndrome Society of the US and the Turner Syndrome Foundation in addition to medical geneticists, pediatricians and pediatric endocrinologists and control participants were recruited by newspaper and internet advertisements. Exclusion criteria for both groups included premature birth (gestational age <34 weeks), low-birth weight (<2000 g), and known diagnosis of a major psychiatric condition (i.e., psychotic or mood disorder) or current neurological disorder, including seizures. This study was approved by the local Institutional Review Board of the Stanford University School of Medicine, and informed written consent was obtained from a legal guardian for all participants, as well as written assent from participants over 7 years of age.

### Neurocognitive data

An extensive neurocognitive assessment was performed, including the Rey–Osterrieth Complex Figure Test (copying of complex figure), 10 core subtests of the WISC-IV (Block Design, Similarities, Digit Span, Picture Concepts, Coding, Vocabulary, Letter-Number Sequencing, Matrix Reasoning, Comprehension, Symbol Search), the WRAVMA Drawing (visual motor), matching (visual-spatial), pegboard (fine-motor), and the NEPSY-II. The following scores of the NEPSY-II were available for all participants: Affect Recognition, Fingertip Tapping (dominant hand, nondominant hand, repetitions, sequences, dominant vs. nondominant hand contrast, repetitions vs sequences contrast), Memory for Faces, Narrative Memory (free and cued recall), Visuomotor precision: total completion time, Visuomotor precision: combined score, Word Generation (semantic total score). For additional 29 subscores of the NEPSY-II containing missing values, we performed data imputation using linear regression (for a complete list of all neurocognitive features see the supplementary material). This resulted in 56 neurocognitive scores for the model. After dimensionality reduction using principal component analysis (PCA), 11 PCs were used as features to build a support vector regression (SVR) model to predict X chromosome mosaicism. Leave-one-out cross validation was performed to test the validity of the model and the generalizability of the outcomes. This procedure was performed by keeping the test and training cases separate. Due to the non-parametric distribution of percentage mosaicism in our sample, Spearman correlation between predicted and actual mosaicism levels was utilized within the cross validation to evaluate the model performance. Cross validation and non-parametric permutation testing were used to test whether the multivariate model significantly predicted the mosaicism level in the test cases.

### Percentage Mosaicism

Mosaicism in all individuals with TS was assessed using standard clinical protocols for fluorescence in situ hybridization (FISH). Peripheral blood was collected from participants and two hundred interphase nuclei were examined using FISH with probes specific for centromeres for the X and Y chromosomes – DXZ1 and DYZ3. For FISH analyses, mosaicism was quantified as the percentage of interphase nuclei demonstrating a single DXZ1 signal, indicating the presence of a single X chromosome, relative to the proportion of nuclei with two DXZ1 signals (indicating presence of two X chromosomes). However, for the purpose of data analysis percentage mosaicism was converted to represent the proportion of 45X cells, e.g. an individual with 75% mosaicism represents a karyotype comprised of 75% 45X cells and 25% 46XX cells. Individuals with any cell types demonstrating a structural abnormality, including ring(X), translocations, deletions, duplications, or any Y chromosome material, were excluded from the study. Therefore, the analysis only included individuals without X chromosome structural abnormalities.

### Structural MRI data

An MRI session was performed at the Stanford University Lucas Center for Medical Imaging on a GE Signa HDxt 3T scanner using an 8-channel receive coil (8HRBRAIN; GE Medical Systems, Milwaukee, WI). All children had the opportunity to familiarize themselves with the scanning procedures during a short training session in a mock scanner, before the start of the MRI scanning session. A high-resolution T1-weighted structural image was acquired using a fast spoiled gradient-echo sequence (FSPGR; 176 slices; repetition time (TR)=8.16; echo time (TE)=3.18; flip angle 12° ; FOV=256×256; voxel size1=mm3; slice thickness=1mm; no slice gap). Quality of the structural images was assessed during the MRI session and the sequence was repeated when needed.

Cortical reconstruction and volumetric segmentation was performed with the Freesurfer image analysis suite (version 5.3), which is documented and freely available for download online (http://surfer.nmr.mgh.harvard.edu/). The technical details of these procedures are described in prior publications (64–77). All structural images were reviewed after FreeSurfer processing and experienced lab members performed manual edits on recon-all outputs to delete dura mater that was segmented incorrectly and to improve white/grey matter segmentation using control points. After manual editing, recon-all processing was repeated before the image was given to an independent expert for final review.

### Support vector regression analysis

Support vector regression (SVR) analyses were performed using our in-house MVPA toolbox on MATLAB (53–55). SVR was implemented using all available features of (1) neurocognitive measures, (2) CV, (3) CT, (4) CSA, and (5) neurocognitive measures and CV to predict the percentage of X chromosome mosaicism. To account for the variability in scaling of the different neurocognitive and neuroanatomical measures, the data was normalized using z-transformed values that were calculated within each measure. Given the high number of independent variables, we performed PCA and the PCs that resulted from eigenvalue-based intrinsic dimensionality reduction were used as features for the SVR. SVR models were trained using a linear kernel. A leave one out cross validation was applied. To evaluate the prediction accuracy by accounting for the non-parametric distribution of percentage mosaicism in our sample, Spearman correlations between predicted and real values were performed. Feature weights of the PC and PC loading to the original features were multiplied for post-hoc estimation of weights for the original set of features to indicate the importance of each feature in their contribution to the SVR. Reconstructed feature weights higher or lower than 1SD from the mean were considered to be of interest and discussed as the features contributing the highest to the SVR predictions.

## Data and code availability

The data of this study is available on: https://osf.io/m4sju/overview?view_only=67246a9a984849b8958a21027c0a9b6c The code of the toolbox used for the SVR analysis is available on: https://code.stanford.edu/cbrain/mvpa-regression

## Author Contributions

DSH conceived and designed the study. IK was responsible for conducting the data analyses and prepared the figures. DSH and ALR oversaw study design and execution. MK oversaw neuropsychological data collection and scoring. IK, MK, LC and TJ scored and analyzed the neuropsychological data. IK analyzed the neuroimaging data. SH contributed to the SVR code development and analytical design. IK, MK and DSH wrote the manuscript, and all authors provided feedback during its preparation.

## Acknowledgments

We would like to thank all the participants and families who participated in the research. We would like to thank Dr. Jennifer Phillips and the students of the Stanford Neurodevelopmental Assessment Practicum Program for their support in neuropsychological assessment administration and scoring, and Sharon Bade Shrestha who assisted with data acquisition.

## References

1. T. T. Mallard, et al., X-chromosome influences on neuroanatomical variation in humans. Nat Neurosci 24, 1216–1224 (2021).

2. P. M. Thompson, et al., The Enhancing NeuroImaging Genetics through Meta-Analysis Consortium: 10 Years of Global Collaborations in Human Brain Mapping. Human Brain Mapping 43, 15–22 (2022).

3. C. Sudlow, et al., UK Biobank: An Open Access Resource for Identifying the Causes of a Wide Range of Complex Diseases of Middle and Old Age. PLOS Medicine 12, e1001779 (2015).

4. A. Raznahan, H. Won, D. C. Glahn, S. Jacquemont, Convergence and Divergence of Rare Genetic Disorders on Brain Phenotypes: A Review. JAMA Psychiatry 79, 818–828 (2022).

5. M. R. Robinson, N. R. Wray, P. M. Visscher, Explaining additional genetic variation in complex traits. Trends in Genetics 30, 124–132 (2014).

6. A. M. Rice, A. McLysaght, Dosage sensitivity is a major determinant of human copy number variant pathogenicity. Nat Commun 8, 14366 (2017).

7. R. A. Polin, A. R. Spitzer, Fetal & Neonatal Secrets (Elsevier Health Sciences, 2013).

8. S. Ohno, Sex chromosomes and sex-linked genes (Springer-Verlag, 1967).

9. P. Avner, E. Heard, X-chromosome inactivation: counting, choice and initiation. Nat Rev Genet 2, 59–67 (2001).

10. C. H. Gravholt, Epidemiological, endocrine and metabolic features in Turner syndrome. European journal of endocrinology 151, 657–688 (2004).

11. B. J. Lahood, G. E. Bacon, Cognitive abilities of adolescent Turner’s syndrome patients. Journal of Adolescent Health Care 6, 358–364 (1985).

12. A. Silbert, P. H. Wolff, J. Lilienthal, Spatial and temporal processing in patients with Turner’s syndrome. Behavior Genetics 7, 11–21 (1977).

13. J. F. Rovet, The psychoeducational characteristics of children with Turner syndrome. Journal of learning disabilities 26, 333–341 (1993).

14. C. M. Temple, Oral fluency and narrative production in children with Turner’s syndrome. Neuropsychologia 40, 1419–1427 (2002).

15. M. M. Murphy, M. M. Mazzocco, Mathematics learning disabilities in girls with fragile X or Turner syndrome during late elementary school. Journal of Learning Disabilities 41, 29–46 (2008).

16. J.-F. Lepage, et al., Genomic Imprinting Effects of the X Chromosome on Brain Morphology. J. Neurosci. 33, 8567–8574 (2013).

17. J.-F. Lepage, P. K. Mazaika, D. S. Hong, M. Raman, A. L. Reiss, Cortical Brain Morphology in Young, Estrogen-Naive, and Adolescent, Estrogen-Treated Girls with Turner Syndrome. Cerebral Cortex 23, 2159–2168 (2013).

18. M. J. Marzelli, F. Hoeft, D. S. Hong, A. L. Reiss, Neuroanatomical spatial patterns in Turner syndrome. NeuroImage 55, 439–447 (2011).

19. C. Rae, et al., Enlarged Temporal Lobes in Turner Syndrome: An X-chromosome Effect? Cerebral Cortex 14, 156–164 (2004).

20. A. Raznahan, et al., Cortical anatomy in human X monosomy. NeuroImage 49, 2915–2923 (2010).

21. S. Xie, et al., The Effects of X Chromosome Loss on Neuroanatomical and Cognitive Phenotypes During Adolescence: a Multi-modal Structural MRI and Diffusion Tensor Imaging Study. Cerebral Cortex 25, 2842–2853 (2015).

22. C. Zhao, G. Gong, Mapping the effect of the X chromosome on the human brain: Neuroimaging evidence from Turner syndrome. Neuroscience & Biobehavioral Reviews 80, 263–275 (2017).

23. D. S. Hong, et al., Influence of the X-Chromosome on Neuroanatomy: Evidence from Turner and Klinefelter Syndromes. J. Neurosci. 34, 3509–3516 (2014).

24. A. R. Zinn, D. C. Page, E. M. C. Fisher, Turner syndrome: the case of the missing sex chromosome. Trends in Genetics 9, 90–93 (1993).

25. X. Deng, et al., Evidence for compensatory upregulation of expressed X-linked genes in mammals, Caenorhabditis elegans and Drosophila melanogaster. Nat Genet 43, 1179–1185 (2011).

26. T. Tukiainen, et al., Landscape of X chromosome inactivation across human tissues. Nature 550, 244–248 (2017).

27. L. G. Biesecker, N. B. Spinner, A genomic view of mosaicism and human disease. Nat Rev Genet 14, 307–320 (2013).

28. D. G. M. Murphy, et al., The effects of sex steroids, and the X chromosome, on female brain function: A study of the neuropsychology of adult turner syndrome. Neuropsychologia 32, 1309–1323 (1994).

29. C. M. Temple, R. A. Carney, Patterns of Spatial Functioning in Turner’s Syndrome. Cortex 31, 109–118 (1995).

30. J. O’Connor, M. Fitzgerald, H. Hoey, The relationship between karyotype and cognitive functioning in Turner syndrome. Ir. j. psychol. med. 17, 82–85 (2000).

31. C. M. Temple, R. A. Carney, Intellectual functioning of children with Turner syndrome: a comparison of behavioural phenotypes. Developmental Medicine & Child Neurology 35, 691–698 (1993).

32. M. W. G. Nijhuis-van der Sanden, P. A. T. M. Eling, B. J. Otten, A review of neuropsychological and motor studies in Turner Syndrome. Neuroscience & Biobehavioral Reviews 27, 329–338 (2003).

33. J. A. Salbenblatt, D. C. Meyers, B. G. Bender, M. G. Linden, A. Robinson, Gross and fine motor development in 45, X and 47, XXX girls. Pediatrics 84, 678–682 (1989).

34. E. Rao, et al., Pseudoautosomal deletions encompassing a novel homeobox gene cause growth failure in idiopathic short stature and Turner syndrome. Nat Genet 16, 54–63 (1997).

35. D. G. M. Murphy, et al., X-chromosome effects on female brain: a magnetic resonance imaging study of Turner’s syndrome. The Lancet 342, 1197–1200 (1993).

36. D. Hong, J. Scaletta Kent, S. Kesler, Cognitive profile of Turner syndrome. Developmental Disabilities Research Reviews 15, 270–278 (2009).

37. J. W. Shaffer, A specific deficit observed in gonadal aplasia (Turner’s syndrome). Journal of Clinical Psychology 18, 403–406 (1962).

38. J. M. Baker, M. Klabunde, B. Jo, T. Green, A. L. Reiss, On the relationship between mathematics and visuospatial processing in Turner syndrome. Journal of Psychiatric Research 121, 135–142 (2020).

39. T. J. Simon, et al., Overlapping numerical cognition impairments in children with chromosome 22q11.2 deletion or Turner syndromes. Neuropsychologia 46, 82–94 (2008).

40. D. H. Skuse, X-linked genes and mental functioning. Human Molecular Genetics 14, R27– R32 (2005).

41. J. Rovet, C. Szekely, M.-N. Hockenberry, Specific arithmetic calculation deficits in children with turner syndrome. Journal of Clinical and Experimental Neuropsychology 16, 820–839 (1994).

42. D. S. Hong, A. L. Reiss, Cognitive and neurological aspects of sex chromosome aneuploidies. The Lancet Neurology 13, 306–318 (2014).

43. F. Printzlau, J. Wolstencroft, D. H. Skuse, Cognitive, behavioral, and neural consequences of sex chromosome aneuploidy. Journal of Neuroscience Research 95, 311–319 (2017).

44. M. M. M. Mazzocco, L. B. Hanich, Math achievement, numerical processing, and executive functions in girls with Turner syndrome: Do girls with Turner syndrome have math learning disability? Learning and Individual Differences 20, 70–81 (2010).

45. T. Green, et al., Effect of sex chromosome number variation on attention-deficit/hyperactivity disorder symptoms, executive function, and processing speed. Developmental Medicine & Child Neurology 64, 331–339 (2022).

46. S. J. Ritchie, et al., Sex Differences in the Adult Human Brain: Evidence from 5216 UK Biobank Participants. Cerebral Cortex 28, 2959–2975 (2018).

47. A. Raznahan, et al., How Does Your Cortex Grow? J. Neurosci. 31, 7174–7177 (2011).

48. D. Beck, et al., Puberty differentially predicts brain maturation in male and female youth: A longitudinal ABCD Study. Developmental Cognitive Neuroscience 61, 101261 (2023).

49. M. A. Tuke, et al., Mosaic Turner syndrome shows reduced penetrance in an adult population study. Genetics in Medicine 21, 877–886 (2019).

50. R. Fernández, J. Méndez, E. Pásaro, Turner syndrome: a study of chromosomal mosaicism. Hum Genet 98, 29–35 (1996).

51. K. R. Held, et al., Mosaicism in 45,X Turner syndrome: does survival in early pregnancy depend on the presence of two sex chromosomes? Hum Genet 88, 288–294 (1992).

52. Accounting for sex in the genome. Nat Med 23, 1243–1243 (2017).

53. S. R. Kesler, et al., Default mode network connectivity distinguishes chemotherapy-treated breast cancer survivors from controls. Proceedings of the National Academy of Sciences 110, 11600–11605 (2013).

54. T. Kim, A. Rahimpour Jounghani, E. Gozdas, S. M. H. Hosseini, Cortical neurite microstructural correlates of time perception in healthy older adults. Heliyon 10, e32534 (2024).

55. E. Gozdas, T. Kim, A. R. Jounghani, S. M. H. Hosseini, Protocol to predict time perception bias from cortical neurite microstructures in healthy older adults. STAR Protocols 6, 103756 (2025).

