## Supplementary Material for "Relationship between X chromosome mosaicism, neuroanatomy and cognitive performance in females"

### Neurocognitive features

The following 56 neurocognitive measures were used as features for the SVR analysis.

REYO:

1. total score copying complex figure

WISC 4:

2. Block Design
3. Similarities
4. Digit Span
5. Picture Concepts
6. Coding
7. Vocabulary
8. Letter-Number Seq.
9. Matrix Reasoning
10. Comprehension
11. Symbol Search

WRAVMA:

12. Drawing (Visual Motor)
13. Matching (Visual-Spatial)
14. Pegboard (Fine-Motor)

NEPSY 2:

15. Affect Recognition
16. Fingertip Tapping-Dominant Hand
17. Fingertip Tapping-Nondominant Hand
18. Fingertip Tapping-Repetitions
19. Fingertip Tapping-Sequences
20. FT-Dominant Hand vs. Nondominant Hand
21. FT Repetitions vs. Sequences
22. Memory for Faces
23. Narrative Memory Free and Cued Recall
24. Narrative Memory Free Recall
25. Visuomotor Precision Total Completion Time
26. Visuomotor Precision
27. Word Generation-Sematic

28. Arrows Total Score – Scaled
29. Auditory Attention Total Correct – Scaled
30. Auditory Attention Combined Scaled Score
31. Response Set Total Correct – Scaled
32. Response Set Combined Scaled Score
33. AA vs. RS Contrast Scaled Score
34. Comprehension of Instructions Total Score – Scaled
35. Imitating Hand Position Total Score – Scaled
36. Inhibition-Naming Completion Time Total – Scaled
37. Inhibition-Inhibition Completion Time Total – Scaled
38. Inhibition-Switching Completion Time Total – Scaled
39. Inhibition Total Errors – Scaled
40. Inhibition-Naming Combined Scaled Score
41. Inhibition-Inhibition Combined Scaled Score
42. Inhibition-Switching Combined Scaled Score
43. IN-Naming vs. Inhibition Contrast Scaled Score
44. IN-Inhibition vs. Switching Contrast Scaled Score
45. List Memory and List Memory Delayed Total Score – Scaled
46. List Memory Learning Effect - Cumulative Percent
47. List Memory Interference Effect - Cumulative Percent
48. List Memory and List Memory Delayed - Delay Effect - Cumulative Percent
49. Memory for Faces Delayed Total Score – Scaled
50. MF vs. MFD Contrast Scaled Score
51. NM Free and Cued Recall vs. Recognition Contrast Scaled Score
52. Picture Puzzles Total Score – Scaled
53. Speeded Naming Total Completion Time – Scaled
54. Speeded Naming Combined Scaled Score
55. Word Generation-Initial Letter Total Score – Scaled
56. WG Semantic vs. Initial Letter Contrast Scaled Score
